# Effect of *Caenorhabditis elegans* U6 Promoters on CRISPR/Cas9-Mediated Gene Editing Efficiency

**DOI:** 10.1101/2025.07.16.664859

**Authors:** Feng Lixiang, Huang Yin, Zhao Rongqian, Zhang Kui, Yang Wenxing

## Abstract

**Objective:** This study investigated the effects of endogenous *C. elegans* U6 promoters on *dpy-10* gene editing efficiency.

**Methods:** We screened endogenous U6 snRNA genes from the WormBase database and constructed 14 editing plasmids targeting *dpy-10* by replacing the U6_*r07e5.16*_ promoter in pSX524 (P_*eft-3*_::*cas9::tbb-2 terminator::U6* _*r07e5.16*_::*dpy-10 sgRNA*) through molecular cloning. Following standardized microinjection protocols, we quantified gene editing efficiency and high-efficiency gene editing index based on *dpy-10* mutant phenotypes in F1 progeny.

**Results:** Fifteen U6 snRNA genes (*r07e5.16, f35c11.9, t20d3.13, k09b11.15, k09b11.16, w05b2.8, c28a5.7, f54c8.9, k09b11.11, k09b11.12, k09b11.14, t20d3.12, f54c8.8, f54c8.10, k09b11.13*) were identified from WormBase database. Comparative analysis revealed four U6 promoters (*w05b2.8, c28a5.7, f54c8.9*, and *k09b11.11*) significantly enhanced gene editing compared to other U6 promoters, including commonly used U6_*r07e5.16*_ and U6_*k09b11.12*_ promoters in *C. elegans* community. Notably, the gRNA^F+E^ scaffold showed no improvement over the gRNA scaffold when paired with the optimal U6_*w05b2.8*_ promoter.

**Conclusion:** This study identifies superior U6 promoters for *C. elegans* gene editing and demonstrates the critical role of promoter optimization in CRISPR systems, providing novel insights for technical refinement.

---

The CRISPR/Cas9 gene editing system has become a widely used tool in the field of biomedicine due to its simple design, operational feasibility, and precise targeting. This system consists of the Cas9 nuclease, which cleaves double-stranded DNA, and a single-guide RNA (sgRNA) that targets specific gene loci. The Cas9 protein can be delivered in the form of in vitro purified protein, expression plasmids, or messenger RNA (mRNA) ^[1-7]^, while the sgRNA can be delivered as sgRNA or via expression plasmids ^[8-10]^. Due to the high cost of synthesizing proteins or RNA, plasmid-based delivery of Cas9 and sgRNA has become the preferred gene editing method for its cost-effectiveness. However, optimizing editing efficiency based on plasmid-mediated gene editing remains a pressing scientific challenge.

As a classic model organism, *Caenorhabditis elegans* is widely used in research areas such as neural regulation, infection and immunity, aging and reproduction, and environmental toxicology mechanisms ^[11-14]^. In *C. elegans* research, the Calarco and Goldstein labs independently developed two Cas9/sgRNA plasmid-based gene editing systems in 2013 ^[1, 2]^. The key differences between these two systems are as follows: the Calarco lab delivered Cas9 and sgRNA via two separate plasmids, while the Goldstein lab used a single plasmid for both; the former employed the promoter of the U6 small nuclear RNA (snRNA) gene U6_*k09b11.12*_ to drive sgRNA transcription, whereas the latter used the U6_*r07e5.16*_ promoter. Although sgRNAs expressed from both U6 promoters have been successfully used for gene editing, reports on their editing efficiency have been inconsistent—some studies suggest the *k09b11.12* promoter is more efficient, while others favor the *r07e5.16* promoter ^[15, 16]^. This indicates that the choice of U6 promoter may influence editing efficiency, but a systematic comparative study is still lacking.

Research by Chen et al. in mammalian cells demonstrated that a modified gRNA scaffold, the flipped plus extended gRNA scaffold (gRNA^F+E^ scaffold), exhibits superior gene editing activity ^[10]^. Additionally, studies have confirmed that the gRNA^F+E^ scaffold significantly enhances gene editing efficiency in *C. elegans* ^[17]^. However, whether the gRNA^F+E^ scaffold can further improve editing efficiency when paired with a more efficient U6 promoter remains to be explored.

Since the knockout or editing of most genes does not result in obvious phenotypic changes, there is no direct screening marker to confirm successful gene editing in the F1 generation of *C. elegans* after using the aforementioned plasmid systems. Consequently, extensive genotyping experiments are required for screening, which increases experimental costs and reduces efficiency. To address this issue, researchers have developed the co-CRISPR strategy ^[18-21]^. This approach involves simultaneously targeting the gene of interest and an easily detectable reference gene, using visible phenotypes (e.g., behavioral or morphological changes) caused by mutations in the reference gene as a screening criterion to improve the efficiency of identifying edits in the target gene. Among co-CRISPR strategies, *dpy-10* is one of the most commonly used reference genes ^[21]^. DPY-10 is a nematode-specific collagen protein, and its *cn64* mutation is a dominant gain-of-function mutation. The molecular mechanism involves the substitution of arginine (Arg92) with cysteine (Cys) at position 92 (p.Arg92Cys) ^[22]^. This mutation disrupts the structural stability of the collagen protein, leading to abnormal cuticle development and three dominant phenotypes ^[18, 22]^: a short and fat body shape (dumpy, DPY, Figure 1), rolling movement (roller, ROL, Figure 1), or both (DPY+ROL). When these phenotypes appear in the F1 generation, it indicates successful gene editing via the co-CRISPR strategy. Subsequently, edits in the target gene can be further screened in the progeny of these mutant individuals.

**Figure 1.**
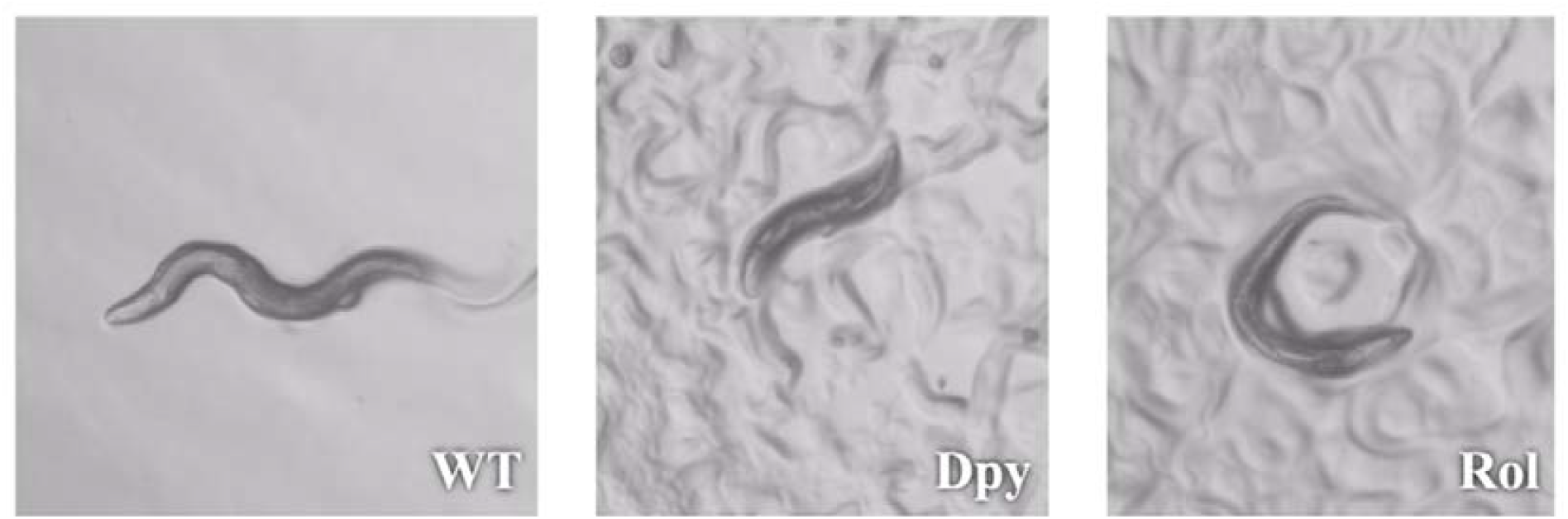
*C. elegans* with wild type, DPY, and ROL phenotypes.

Based on this research background, this study will compare the effects of 15 endogenous *C. elegans* U6 snRNA gene promoters and the gRNA^F+E^ scaffold on *dpy-10* gene editing efficiency, aiming to provide a theoretical basis for optimizing the CRISPR/Cas9 gene editing system in *C. elegans*.

## 1 Materials and Methods

### 1.1 Experimental Animals and Reagents

Wild-type N2 *Caenorhabditis elegans* hermaphrodites were purchased from the Caenorhabditis Genetics Center (CGC, USA). All worms were maintained on nematode growth medium (NGM) plates and routinely cultured with *Escherichia coli* OP50 [^11, 23]^. The 2× high-fidelity PCR premix (Cat. No.: HY-K0533MCE) was purchased from MCE Biotech, and the seamless cloning kit (Cat. No.: MC40101M) was obtained from Monad Biotech.

### 1.2 Experimental Methods

#### 1.2.1 Plasmid Construction Strategy

All targeting plasmids were constructed using pDD162 (P_*eft-3*_::Cas9 + Empty sgRNA) as the template ^[1]^. The pDD162 plasmid contains the P_*eft-3*_::Cas9 expression cassette, the U6_*r07e5.16*_ promoter (derived from the *r07e5.16* gene), and an sgRNA scaffold lacking target-complementary sequences. The complementary sequences were replaced with a 20-bp targeting sequence for the *dpy-10* gene (Figure 2) via molecular cloning, resulting in the pSX524 plasmid. The pSX524 plasmid used in this study was kindly provided by Prof. Wei Zou from Zhejiang University. The U6_*r07e5.16*_ promoter in the original vector was replaced with other U6 gene promoters using seamless cloning to generate the series of targeting plasmids used in this study. All recombinant plasmids were verified by DNA sequencing. Detailed molecular cloning protocols, complete plasmid DNA sequences, and primer sequences used in plasmid construction are provided in Supplementary Tables 1 and 2 (Figshare.com ^[24]^).

**Table 1.** U6 snRNA-related information.

| U6 snRNA | alternative name | plasmid | length of U6 promoter (bp) | genomic location | the number of microinjections |
| --- | --- | --- | --- | --- | --- |
| F35C11.9 | U6 | YW611 | 500 | II:0.75 +/- 0.000 cM | 8 |
| F54C8.8 | U6-1 | YW622 | 500 | III:0.52 +/- 0.000 cM | 14 |
| F54C8.9 | U6-2 | YW617 | 500 | III:0.52 +/- 0.000 cM | 9 |
| F54C8.10 | U6-3 | YW623 | 500 | III:0.52 +/- 0.000 cM | 14 |
| C28A5.7 | U6-4 | YW616 | 500 | III:-3.17 +/- 0.000 cM | 42 |
| K09B11.11 | U6-8 | YW618 | 500 | IV:9.25 +/- 0.000 cM | 14 |
| K09B11.12 | U6-9 | YW619 | 500 | IV:9.25 +/- 0.000 cM | 16 |
| K09B11.13 | U6-10 | YW624 | 500 | IV:9.26 +/- 0.000 cM | 7 |
| K09B11.14 | U6-11 | YW620 | 500 | IV:9.26 +/- 0.000 cM | 7 |
| K09B11.15 | U6-12 | YW613 | 500 | IV:9.26 +/- 0.000 cM | 8 |
| K09B11.16 | U6-13 | YW614 | 500 | IV:9.26 +/- 0.000 cM | 6 |
| R07E5.16 | U6-15 | SX524 | 350 | III:-3.17 +/- 0.000 cM | 16 |
| T20D3.12 | U6-17 | YW621 | 500 | IV:4.18 +/- 0.000 cM | 7 |
| T20D3.13 | U6-18 | YW612 | 500 | IV:4.18 +/- 0.000 cM | 12 |
| W05B2.8 | U6-19 | YW615 | 500 | III:5.82 +/- 0.000 cM | 11 |
|  |  |  |  |  | Total: 191 |

**Figure 2.**
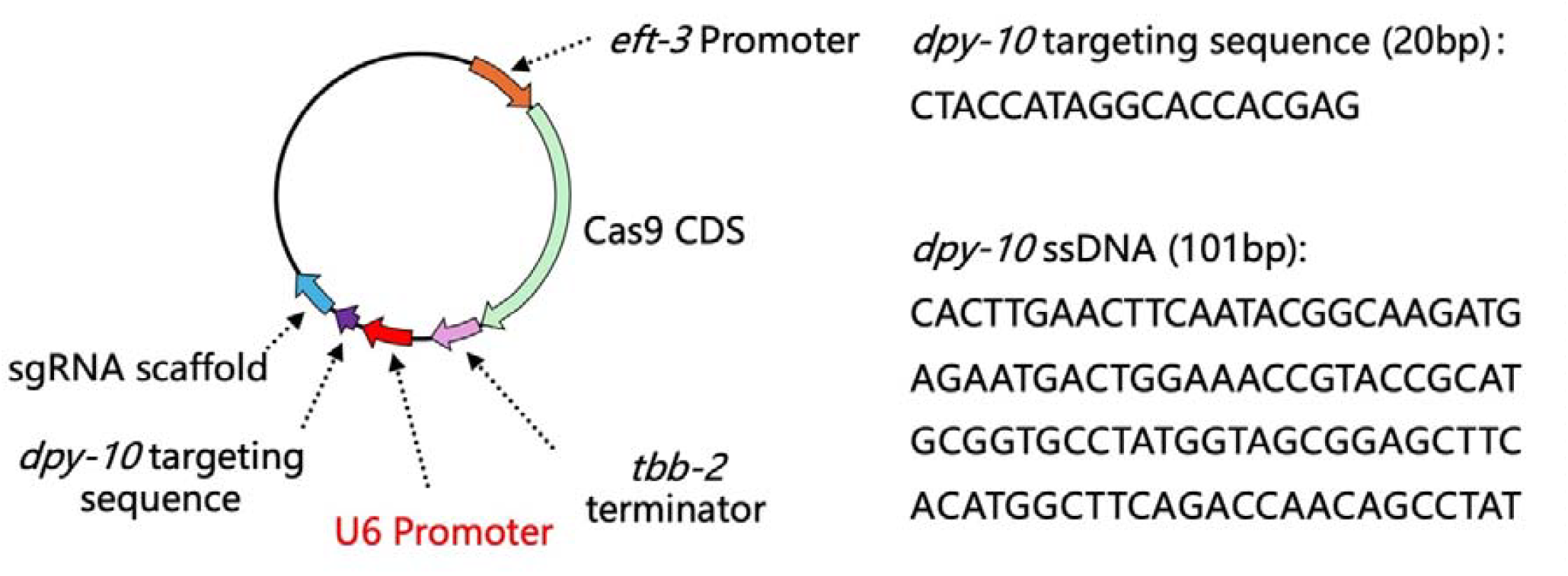
Composition of U6 promoter-related plasmids.

#### 1.2.2 Microinjection Protocol

The *C. elegans* gonad consists of two symmetrical U-shaped arms. To enhance experimental feasibility and minimize variability, microinjection was performed to deliver the targeting plasmids into the distal gonadal region near the head. The plasmid mixture for injection was prepared in a 10 μL total volume, containing the targeting plasmid (30 ng/μL), *dpy-10* gene editing template single-stranded DNA (ssDNA, 101 bp, sequence shown in Figure 2, 0.6 μmol/L), a microinjection marker plasmid (P_*unc-122*_::GFP, 30 ng/μL), and a concentration-balancing plasmid (pUC19, 30 ng/μL), with the final volume adjusted to 10 μL using double-distilled water. Each worm was injected with 20 nL of the mixture. The pYW616 plasmid mixture was used as the standard control. To ensure experimental consistency, each microinjection session (regardless of the number of plasmid mixtures tested) included a pYW616 plasmid mixture as an internal reference.

#### 1.2.3 Quantification of Gene Editing Efficiency

To quantitatively assess gene editing efficiency after microinjection, each successfully injected P0 worm was individually cultured, and plates were replaced daily to collect F1 progeny. When F1 worms reached the L4 larval stage or young adult stage, the total number of worms and those exhibiting *dpy-10* mutant phenotypes were counted under a stereomicroscope. Based on these data, two quantitative metrics were established: gene-editing efficiency and the high-efficiency gene-editing index (HEGE Index). Gene-editing efficiency was defined as the percentage of F1 progeny from a single P0 worm that displayed *dpy-10* mutant phenotypes. A gene-editing efficiency ≥20% was classified as a high-efficiency editing event, and the HEGE Index was calculated by multiplying the proportion of high-efficiency editing events for a given plasmid mixture by the actual gene-editing efficiency of the corresponding P0 worms. This index reflects both the success rate and the reliability of high-efficiency editing.

### 1.3 Statistical Analysis

Data are presented as box plots, showing the median, interquartile range, maximum, and minimum values, with “+” indicating the mean. Statistical analyses were performed using GraphPad Prism 10. Kruskal-Wallis tests followed by Dunn’s multiple comparisons were used for Figures 3 and 4, while the Mann-Whitney test was applied for Figure 5. A *P*-value <0.05 was considered statistically significant.

**Figure 3.**
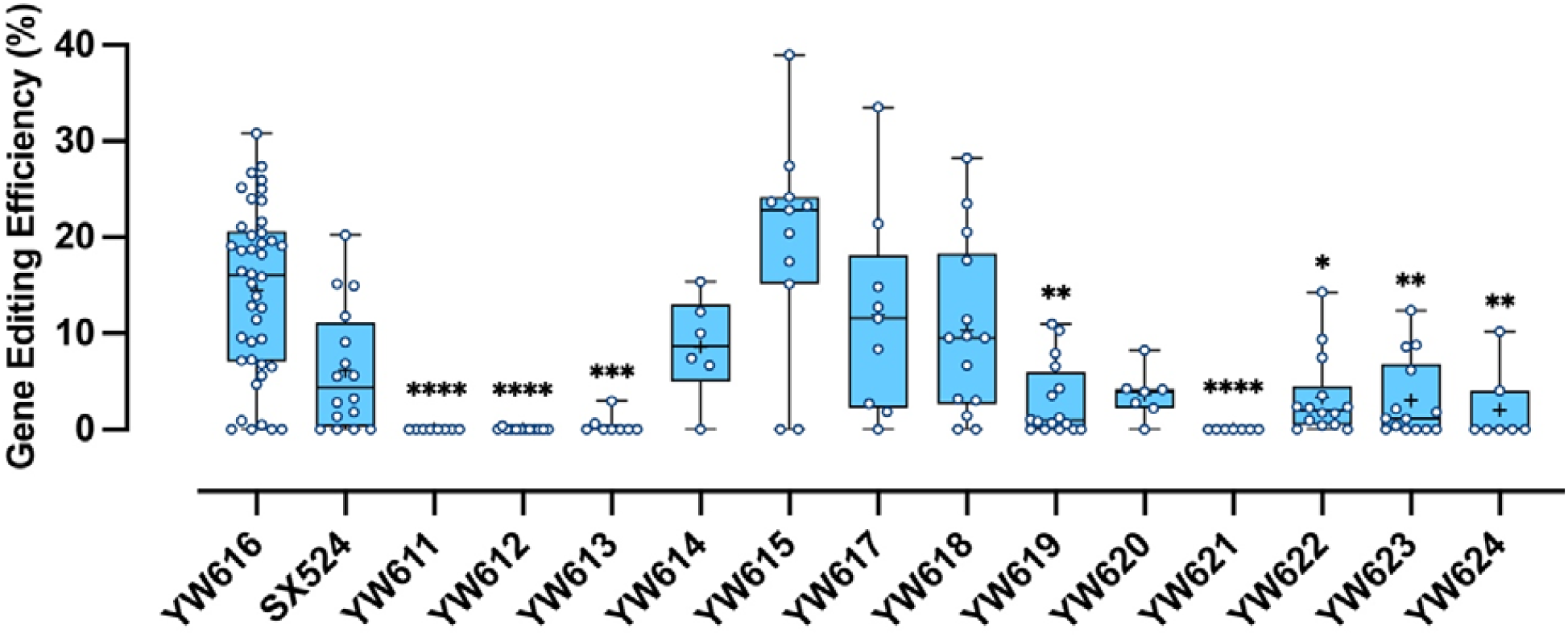
Impact of U6 promoter on gene-editing efficiency. ^****^*P*<0.0001, ^***^*P*<0.001, ^**^P<0.01, ^*^*P*<0.05. vs YW616. From left to right, the sample sizes (n) are 42, 16, 8, 12, 8, 6, 11, 9, 14, 16, 7, 7, 14, 14, 7, respectively.

**Figure 4.**
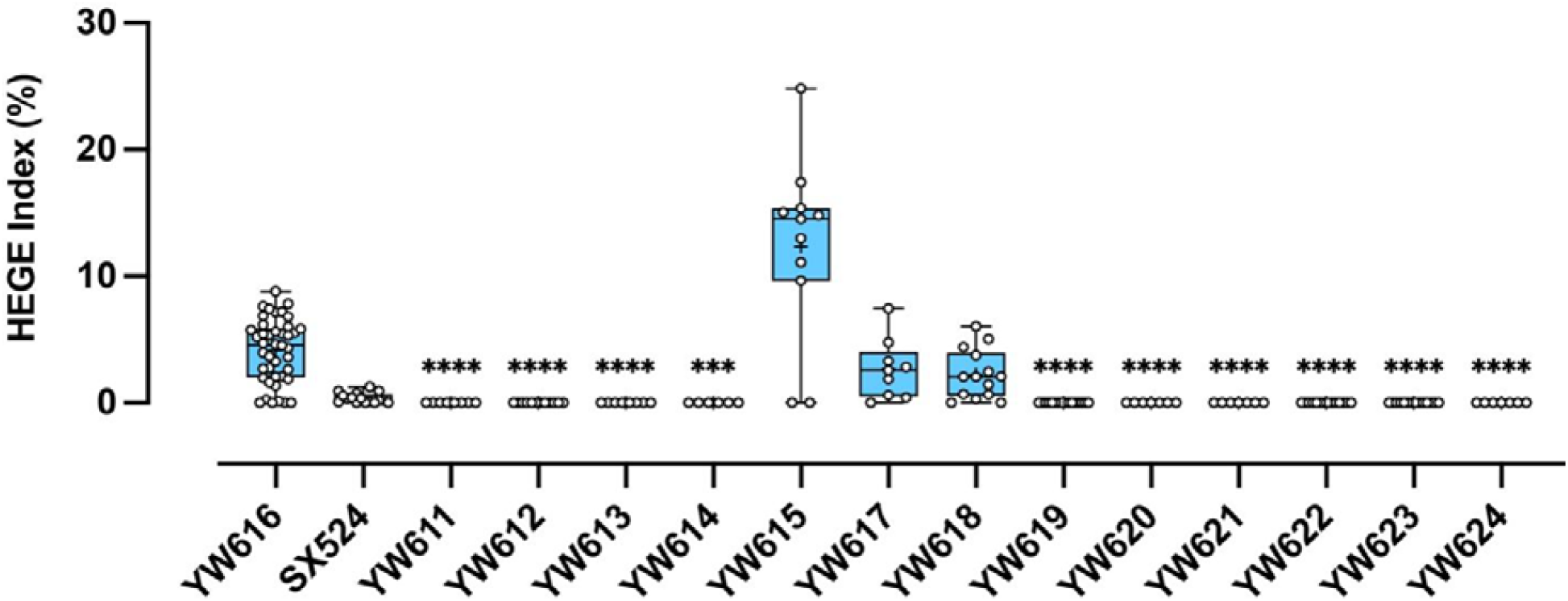
Impact of U6 promoter on high efficiency gene-editing index. ^****^*P*<0.0001, ^***^*P*<0.001, vs YW616. From left to right, the sample sizes (n) are 42, 16, 8, 12, 8, 6, 11, 9, 14, 16, 7, 7, 14, 14, 7, respectively.

**Figure 5.**
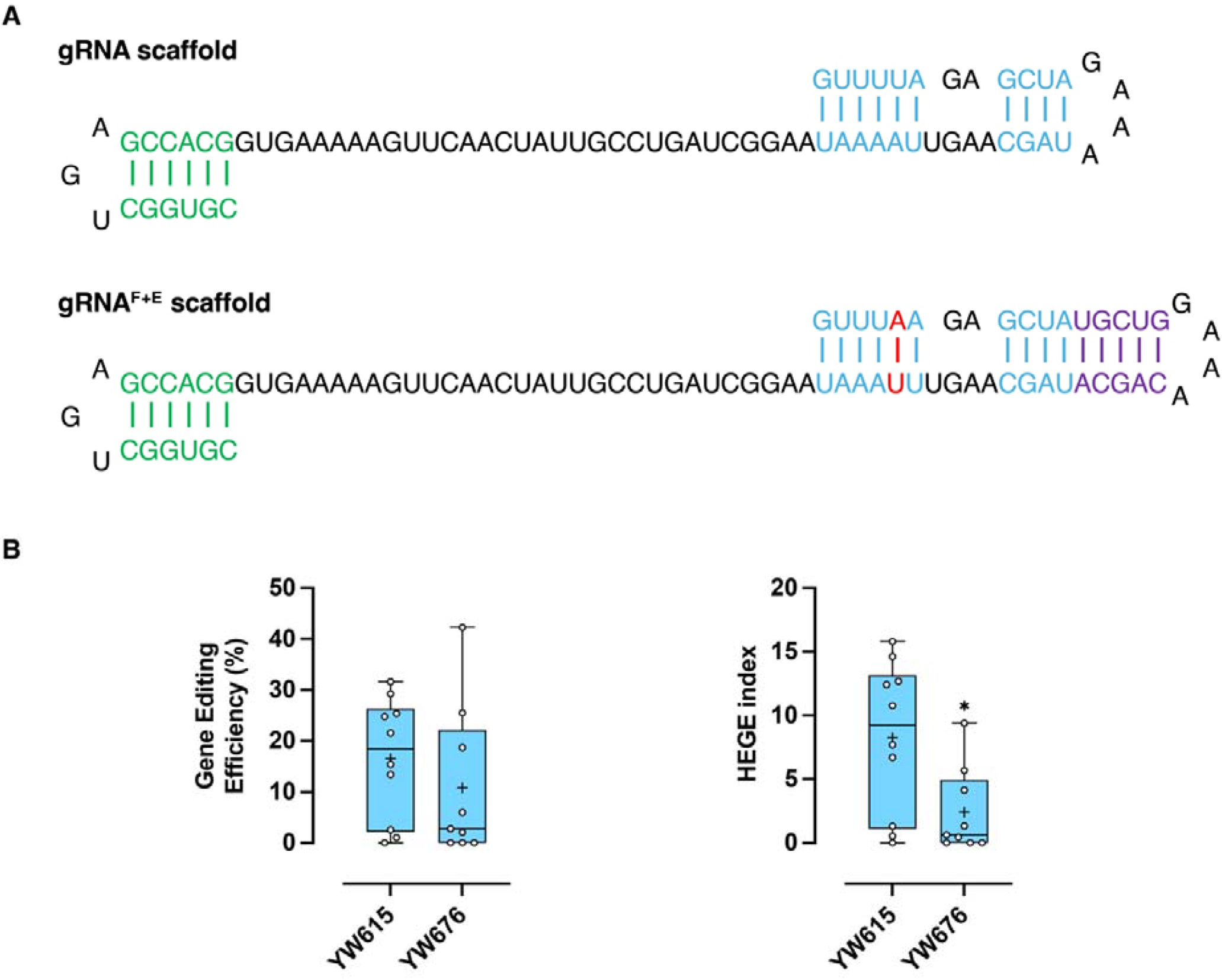
Impact of gRNA scaffold on gene-editing. ^*^*P*<0.05; vs YW615. n = 9-10.

## 2 Results

### 2.1 Construction of Plasmid Vectors with Different U6 Promoters

By searching the “Gene” option in the WormBase database (www.wormbase.org) using the keyword “U6”, we identified 15 U6 snRNA genes (Table 1). The pSX524 plasmid (*Peft-3::Cas9::tbb-2 terminator::U6r07e5.16::dpy-10 sgRNA*) contains the Cas9 gene under the control of the P_*eft-3*_ promoter, with the 3’ untranslated region (UTR) of the *tbb-2* gene serving as the transcriptional terminator. Downstream of the *Cas9* expression cassette is the *dpy-10* sgRNA expression cassette driven by the U6_*r07e5.16*_ promoter (Figure 2). In this study, we constructed the pYW611–pYW624 plasmids by replacing the U6_*r07e5.16*_ promoter in pSX524 with the other 14 U6 promoters. The correspondence between plasmid names and U6 promoters is summarized in Table 1.

### 2.2 Different U6 Promoters Lead to Varying Gene Editing Efficiencies

The *dpy-10* cn64 mutation disrupts the structural stability of collagen, leading to abnormal cuticle development and three dominant phenotypes [18, 22]: dumpy (dpy), roller (rol), and dumpy+roller (dpy+rol) (Figure 1). By using sgRNA to guide Cas9 to target specific genomic loci, along with a 101-bp single-stranded DNA (ssDNA, Figure 2) as a DNA repair template, wild-type *C. elegans* subjected to gene editing will produce progeny carrying the cn64 mutation. These mutant phenotypes can be identified under a stereomicroscope. If wild-type P0 worms are microinjected for gene editing, the F1 generation will exhibit the aforementioned cn64 mutant phenotypes. Therefore, the proportion of phenotype-positive F1 progeny serves as a reliable metric for assessing gene editing efficiency.

To evaluate the impact of 15 different U6 promoters on gene editing efficiency, we conducted 191 microinjection experiments using the pSX524 plasmid and 14 derived plasmids (pYW611–pYW624) (Table 1). The results showed that four plasmids—pYW611, pYW612, pYW613, and pYW621—exhibited near-zero editing efficiency, indicating that their respective U6 snRNA gene promoters (*f35c11.9, t20d3.13, k09b11.15, t20d3.12*) were nearly incapable of driving effective gene editing (Figure 3).

The pYW616 group demonstrated significantly higher editing efficiency than the pYW619 group (*P* < 0.01), suggesting that the *c28a5.7* promoter outperforms the *k09b11.12* promoter used by the Calarco lab. Compared to the *r07e5.16* promoter (pSX524 group) employed by the Goldstein lab, the pYW616 group showed a trend toward higher editing efficiency, though the difference was not statistically significant (*P* > 0.05), indicating that the *c28a5.7* promoter did not significantly surpass *r07e5.16*. Additionally, no significant differences were observed between the pYW616 group and the pYW614, pYW615, pYW617, pYW618, or pYW620 groups (*P* > 0.05), suggesting that the *k09b11.16, w05b2.8, f54c8.9, k09b11.11*, and *k09b11.14* promoters performed similarly to *c28a5.7*. In contrast, the pYW616 group exhibited significantly higher editing efficiency than the pYW621 (*P* < 0.0001), pYW622 (*P* < 0.05), pYW623 (*P* < 0.01), and pYW624 (*P* < 0.01) groups, demonstrating that the *c28a5.7* promoter outperformed the *t20d3.12, f54c8.8, f54c8.10*, and *k09b11.13* promoters.

### 2.3 The *c28a5.7, w05b2.8, f54c8.9*, and *k09b11.11* Promoters Exhibit Higher HEGE Indices

As shown in Figure 3, some individuals in each experimental group displayed zero editing efficiency, indicating that even with successful microinjection, gene editing events remain probabilistic and not guaranteed. Given the stochastic nature of gene editing, we further quantified editing performance using the HEGE index.

As illustrated in Figure 4, the HEGE index of the pYW616 group was significantly higher than those of the pYW611–pYW614 and pYW619–pYW624 groups (*P* < 0.001 or *P* < 0.0001) and showed a trend toward surpassing the pSX524 group (*P* = 0.0512). Meanwhile, no significant differences were observed among the pYW615–pYW618 groups (*P* > 0.05). These results demonstrate that the *c28a5.7, w05b2.8, f54c8.9*, and *k09b11.11* promoters significantly enhance the HEGE index, making them superior choices for HEGE in *C. elegans*.

### 2.4 No Significant Impact of Two Different gRNA Scaffolds on Gene Editing Mediated by the *w05b2.8* Promoter

In this study, all plasmids employed the conventional gRNA scaffold. To investigate whether the gRNA^F+E^ scaffold (Figure 5A) could further enhance editing efficiency when combined with high-performance U6 promoters, we conducted comparative experiments to evaluate the effects of these two scaffolds. Figures 3 and 4 demonstrate that among the four U6 promoter plasmids showing higher editing efficiency, YW615 exhibited the highest median values for both gene editing efficiency and HEGE Index, suggesting it might be the most effective among these high-performance U6 promoters. Based on this finding, we constructed the YW676 plasmid using YW615 as the backbone, with the sole difference being the replacement of the conventional gRNA scaffold with the gRNA^F+E^ scaffold. As shown in Figure 5B, no statistically significant difference was observed in gene editing efficiency between the two gRNA scaffolds (P>0.05). However, the YW676 group using the gRNA^F+E^ scaffold showed a significantly reduced HEGE Index (P<0.05). This reduction may be attributed to a higher probability of failed editing events when using the gRNA^F+E^ scaffold, consequently decreasing the overall HEGE index. These results indicate that while the gRNA^F+E^ scaffold has been reported to enhance editing efficiency in other systems, its benefits may not extend universally to all experimental conditions, particularly when combined with already highly efficient U6 promoters in *C. elegans*. The observed decrease in HEGE Index suggests that scaffold optimization should be carefully evaluated in the context of specific experimental setups and promoter combinations.

## 3 Discussion

*Caenorhabditis elegans* has become a crucial model organism for gene function studies due to its simple cultivation and ease of genetic manipulation. By introducing precise targeted mutations into its endogenous genes, researchers can deeply investigate gene-phenotype relationships. Since the first application of the CRISPR-Cas9 system in *C. elegans* in 2012, this technology has significantly advanced nematode genetics ^[25]^. Although optimization efforts for CRISPR-Cas9 in *C. elegans* have continued over the past decade, editing efficiency remains suboptimal.

Delivering Cas9 protein and sgRNA via plasmid injection is an economical and straightforward gene editing strategy. In this study, we compared the effects of 15 different U6 promoters on editing efficiency and found that the *w05b2.8, c28a5.7, f54c8.9*, and *k09b11.11* promoters exhibited higher editing efficiency and success rates, significantly outperforming the widely used *k09b11.12* (from the Calarco lab) and *r07e5.16* (from the Goldstein lab) promoters. Although the *r07e5.16* promoter showed a trend toward higher editing efficiency than *k09b11.12*, the difference was not statistically significant. Figure 3 reveals that most data points for the *k09b11.12* promoter were close to zero, indicating its extremely low efficiency in inducing gene editing in *C. elegans*. Therefore, between these two promoters, we still recommend prioritizing *r07e5.16*.

Froehlich et al. compared the editing efficiency of the *w05b2.8* and *r07e5.16* promoters and found that *w05b2.8* performed better ^[26]^, consistent with our results. They observed that *w05b2.8* drove higher expression in the gonad than *r07e5.16*, suggesting this as a potential mechanism for its superior efficiency. In our study, except for the pYW676 plasmid, the only variable among constructs was the U6 promoter used. Thus, we hypothesize that differences in editing efficiency primarily stem from variations in sgRNA expression levels driven by these promoters in the gonad. Specifically, the *c28a5.7, w05b2.8, f54c8.9*, and *k09b11.11* promoters likely mediate higher sgRNA transcription in gonadal tissue. In summary, we recommend prioritizing *w05b2.8, c28a5.7, f54c8.9*, and *k09b11.11* promoters for HEGE via enhanced snRNA expression in the gonad.

Compared to the original gRNA scaffold, the gRNA^F+E^ scaffold eliminates potential RNA polymerase III terminators through A-U base pair substitutions and extends the Cas9-binding domain ^[10]^. Previous studies have shown that the gRNA^F+E^ scaffold can improve editing efficiency in various model organisms ^[10, 17]^. However, in our study, when paired with high-efficiency U6 promoters, the gRNA^F+E^ scaffold did not outperform the conventional gRNA scaffold. Our findings suggest that optimizing U6 promoter-driven sgRNA transcription in the gonad may be more effective for enhancing editing efficiency than modifying the gRNA scaffold alone. This provides new insights for optimizing plasmid-based gene editing strategies.

## Author Contribution

FENG Lixiang: Data collection and analysis, drafting the initial manuscript. HUANG Ying and ZHAO Rongqian: Data validation, manuscript revision. ZHANG Kui: Experimental design and conceptualization, research project management. YANG Wenxing: Microinjection, funding acquisition, experimental design and conceptualization, manuscript revision and finalization. All authors have approved the final version of the manuscript and agree to be accountable for all aspects of the work.

